# Immature Myeloid Cell Deposition in Old Bone Marrow Revealed by Single-Cell Transcriptome Analysis

**DOI:** 10.1101/2022.10.15.512392

**Authors:** Woo Jin Kim, Ki-Tae Kim, Jae-I Moon, Seung Gwa Park, Young-Dan Cho, Hyun Jung Kim, Hye-Rim Shin, Heein Yoon, Hyun-Mo Ryoo

**Affiliations:** Department of Molecular Genetics & Dental Pharmacology, School of Dentistry and Dental Research Institute, Seoul National University, Seoul, South Korea; Epigenetic Regulation of Aged Skeleto-Muscular System Laboratory, School of Dentistry and Dental Research Institute, Seoul National University, Seoul, South Korea; Department of Periodontology, School of Dentistry and Dental Research Institute, Seoul National University, Seoul, South Korea

**Author notes:** These authors contributed equally (Woo Jin Kim, Ki-Tae Kim).

## Abstract

Aging causes dysfunction of innate immunity, although hematopoietic stem cells of aged bone marrow (BM) show an increased differentiation potential to myeloid lineage cells. The alteration of cellular heterogeneity and intercellular communications between BM immune cells may provide important clues to understanding age-dependent immune dysfunction. Here, we provide a deep single-cell transcriptomic analysis of total immune cell populations of young and old BM. We identified the well-organized differentiation status of 11 myeloid/lymphoid lineage cell populations and age-dependent alterations in the proportions of cells. The neutrophil lineages showed the most prominent alteration by aging, and subclustering of neutrophils indicated that the specific immature neutrophils are increased in old BM. In addition, we identified age-dependent alterations in secretory phenotypes associated with a decline in innate immunity and immune cell differentiation. Among these secretory phenotypes, SPP1 could be suggested as a representative signal that triggers myeloid skewing and immature neutrophil deposition in aged BM. Collectively, these results provide a novel link between the altered immune cell proportions in BM and age-dependent dysregulation of innate immunity.

## Introduction

Aging is a multimodal process involving diverse organs, tissues and cells that undergo progressive degenerative changes; thus, bulk cell- or tissue-level approaches to address this complexity are limited^1^. It is well known that systemic inflammation and dysregulation of the immune response are hallmarks of aging, but not all immune processes are uniformly sensitive to aging, suggesting that the diversity of immune cell populations is substantially altered in elderly individuals^2^.

The effects of aging on the innate immune system have not been well studied, in contrast to those on adaptive immunity, until recently^3^. The diverse immune cell lineages in the bone marrow (BM) can be precursors for the generation of innate and adaptive immune cells and show heterogeneous aging phenotypes that reflect different developmental, differentiation, and activation contexts^4^. Previously, many studies in elderly individuals have shown that dysregulation of the immune system is manifested by uncontrolled systemic inflammation, often referred to as ‘inflammaging’^2^. This aging-associated dysregulation results in elevated levels of basal inflammation and an associated impaired ability to elicit an efficient immune response to pathogens or vaccinations^5^.

Recent single-cell RNA sequencing (scRNA-seq) studies have highlighted numerous transcriptomic changes at the single-cell level in various organs^6,7^. Additionally, deep scRNA-seq analyses have revealed substantial tissue-specific heterogeneity among myeloid cells, connected with age-dependent pathophysiological changes in lung, peritoneum, liver, spleen and mammary tissue^8–10^. Although the BM has a key role in the innate immune response, the age-dependent alteration of cellular heterogeneity and intercellular communication between various immune cells in the BM is not completely understood. Thus, we aimed to evaluate unique immature myeloid deposition and identify linked subpopulation changes in response to age-dependent dysregulation of innate immunity using deep scRNA-seq analysis of young, middle-aged and old BM. Computational prediction of intercellular communication using CellChat and NicheNet was used to identify 11 significant signaling pathways responding to alterations in ligand–receptor profiles. Representatively, SPP1, known to be a key regulatory factor of myeloid/lymphoid population balance and to be depleted in old BM^11^, was suggested to be exclusively expressed in young mature neutrophils, and elevated levels of immature myeloid cells in old BM contributed to SPP1 depletion in old BM. These results provide a link between the alterations in immune cell proportions and age-dependent dysregulation of innate immunity, as well as a systemic connection underlying multiorgan aging.

## Results

### Immune cell atlas of bone marrow in aging

To study the changes in the cellular landscape of BM tissue upon aging, we performed scRNA-seq using a microfluidic droplet platform on BM from young (3 months old), middle-aged (12 months old) and old (20 months old) male mice **(Figure 1A)**, which correspond to human early adulthood (20-30 years old), middle age (38-47 years old) and old age (56-69 years old), respectively^12^.

**Figure 1.**
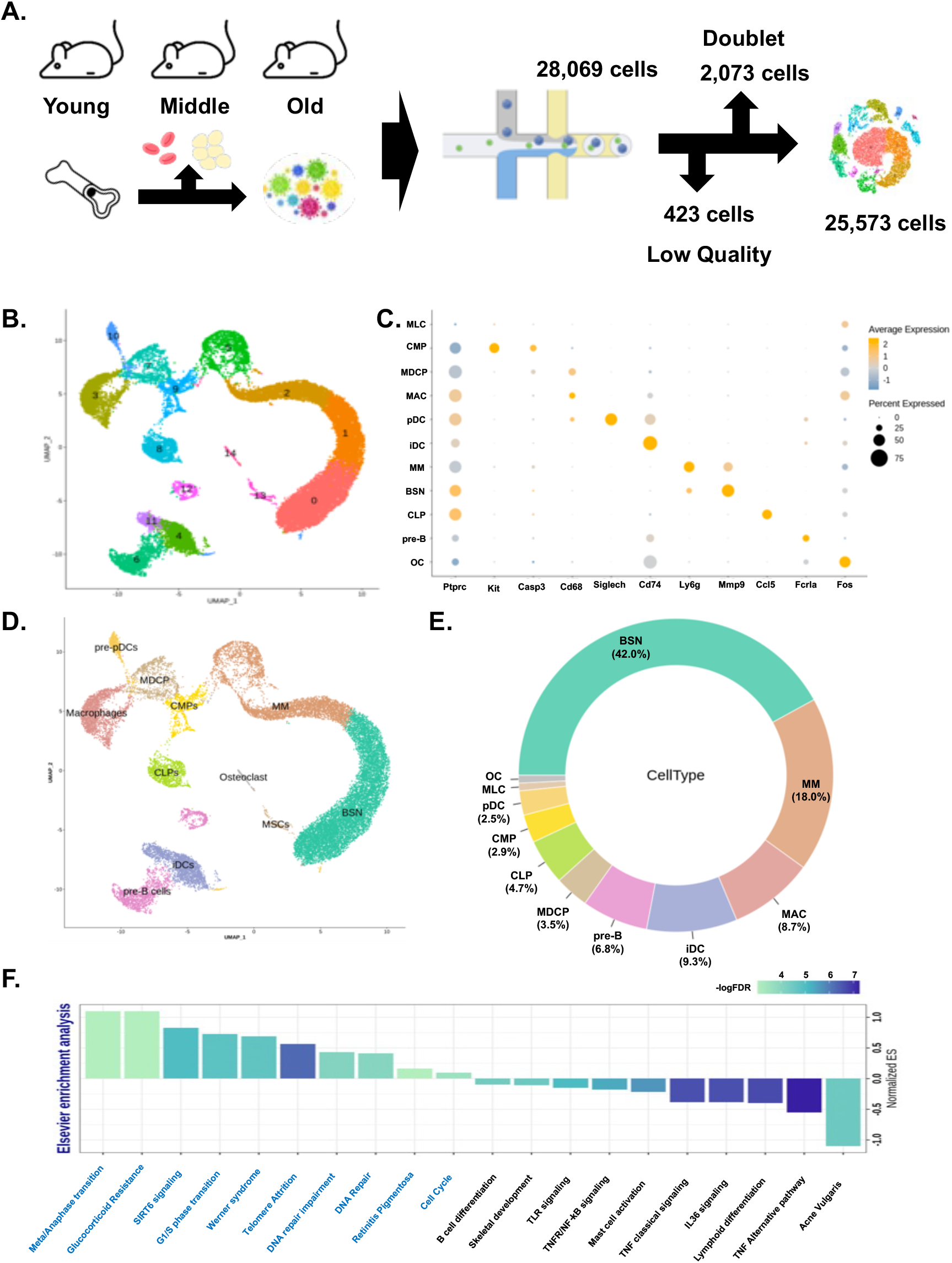
Immune cell atlas of BM in aging revealed by scRNA-seq. **A.** Schematic of the experimental design for scRNA-seq. BM cells were collected from young (3 months of age), middle-aged (12 months of age) and old (20 months of age) mouse femurs and then processed by scRNA-seq using the microfluidic platform. (N=3) **B.** UMAP of the total BM immune cell cluster. **C.** Expression of canonical markers in 11-cell types. The color code indicates the expression level, and the bubble chart indicates the proportion of cells. **D.** Cell-type annotation results identified 11 cell clusters. **E.** Circular plot describing the proportion of cell types. **F.** Enrichment analysis of differentially expressed genes in old BM versus young BM. The Y-axis indicates normalized ES, and the color code indicates FDR. Abbreviations: BSN (banded and segmented neutrophil), MM (metamyelocyte), MAC (macrophage), iDC (immature dendritic cell), pre-B (B-cell precursor), MDCP (macrophage/dendritic cell progenitor), CLP (common lymphoid precursor), CMP (common myeloid precursor), pDC (plasmacytoid dendritic cell), MLC (mesenchymal lineage cell), OC (osteoclast precursor).

To obtain total immune cells, we removed red blood cells and adipocytes from the bone marrow tissue. After stringent filtering of cells to eliminate low-quality outputs and potential doublets, we captured a total of 25,573 cells and 32,285 genes. Single cells were clustered based on gene expression profiles using Seurat, and the clusters were visualized using uniform manifold approximation and projection (UMAP)^13^. Division of the initial cell clusters using principal component analysis of the most variable genes between cells, followed by Louvain and Leiden graph-based clustering, identified 15 well-distinguished clusters **(Figure 1B and Supplementary Figure 1)**.

To define cell types and map clusters, we applied a greedy algorithm with a maximum frequency of 2 databases, CellMarker and the Mouse Cell Atlas, and confirmed the uniformity of the clusters using canonical markers.^14,15^ As expected, we detected immature myeloid lineage cells, such as common myeloid progenitor cells (CMPs) expressing *c-kit*, CD34, CD45 and *Sca-1*, and metamyelocytes (MM), expressing CD11b, *Cxcr4*, *Gr-1*, *Camp*, *Ceacam1*, *Ltf*, *Olfm4*, and *Orm1* and negative for CD115. Mature bone marrow neutrophils are usually banded or segmented nucleus neutrophils (BSNs) and express *Adam8*, *Atp11a*, *Ctsd*, *Fcer1g*, *Lamp2*, *Mgam*, *Plaur*, *Slc11a1*, *Timp2* and matrix metalloproteinases (MMPs), such as *Mmp8* and *Mmp9*; they also express TNF (tumor necrosis factor)-related genes, such as *Tnfaip6* and *Tnfsf14*, and MM marker genes, such as CD11b and *Gr-1*.

Monocyte-derived cell lineages include macrophage/dendritic cell progenitors (MDCPs), dendritic lineages such as immature dendritic cells (iDCs) (characterized by CD74, CD86, *Cst3*, *H2-Ab1* and *H2-Eb1* expression) and immature plasmacytoid dendritic cells (pre-pDCs) (significantly expressing *Siglech*^16^), macrophages (MACs) (characterized by CD68, *Itgam*, CD80, *Fcgr1*, *Fcgr3*, *Nos2*, *Gpr18*, *Fpr2* and CD38 expression) and preosteoclasts (OCs) (high expression of *c-Fos*^17^).

Lymphoid lineage cells were rarely detected, and the representative lymphoid progenitor cells (CLPs) were enriched for key markers^18^, such as CCL5, CD10, CD38, CD52, CD62, and *Csf3R*, with high expression of CD45; immature B cells (pre-B) were distinguished by *Fcrla* and *Vpreb1* expression^19^ **(Figure 1C)**.

As a result of cluster mapping, the 15 clusters were condensed into 11 clusters according to similar cell-type marker expression **(Figure 1D)**. In these data, all myeloid lineages were classified into 8 subtypes representing 87.7% of cells. Major clusters 0 and 1 were combined as BSNs (42% of total cells), and clusters 2 and 5 were merged as MMs (18% of total cells). Other myeloid lineages, namely, CMPs (2.9%), MDCPs (3.5%), MACs (8.7%) and pre-pDCs (2.5%), formed a minor proportion of the cells. All lymphoid lineages comprised 11.4% of the cells, and mesenchymal lineages (MLC) contributed less than 1% **(Figure 1E)**.

Differential expression analysis identified 123 upregulated genes and 36 downregulated genes in old BM **(Supplementary Figure 2)**. Overrepresentation analysis performed to identify biological pathways in those genes indicated that they fell primarily into four functional categories **(Figure 1F)**. More specifically, upregulated genes in old BM were associated with the cell cycle (Cdc20, Pinx1), SIRT signaling (Sirt6, Xrcc6) and DNA damage repair (Bbc3, Atr). An analysis of the genes downregulated in old BM showed that the innate immune reaction was dependent on TNF signaling (associated with Il1a, Bcl2l1 and Tgm2), TLR (Toll-like receptor) signaling and lymphoid lineage differentiation.

These results suggest significant changes in the number and function of myeloid cells, which account for a high proportion of the old BM. Therefore, we further analyzed the nature of the myeloid predisposition of old BM cells.

### Myeloid lineage skewing of old BM immune cells

To characterize the transcriptional dynamics of the temporal process of cell differentiation in myeloid and lymphoid lineages, we used an unsupervised algorithm that increases the temporal resolution of transcriptome dynamics collected at multiple time points in Monocle 3^20^.

We identified lineage changes in the expression of previously known genes associated with immune cell differentiation over kinetic time (pseudotime) across all cells. In view of the myeloid and lymphoid cells defined by Louvain clustering as subtrajectories, the myeloid lineages clearly represented branching trajectories originating from CMPs, and the functional maturation of neutrophils was a significant direction over increasing pseudotime. However, we could not observe a clear dissociation of the lymphoid lineages in the total trajectory **(Figure 2A)**. To define lymphoid differentiation, we mapped lymphoid lineages separately, which showed branched pre-T- and B-cell differentiation trajectories **(Figure 2B)**.

**Figure 2.**
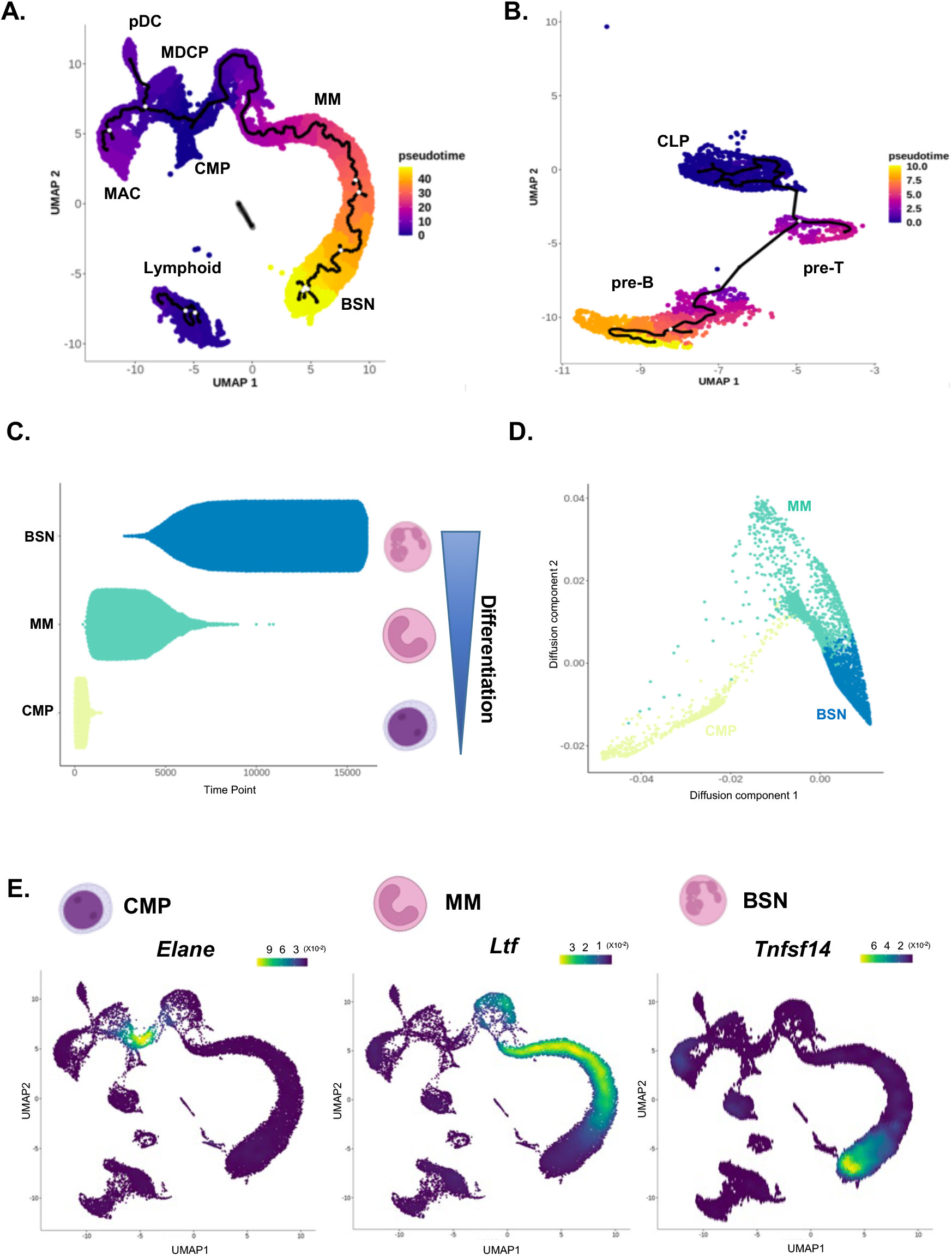
Trajectory analysis of myeloid lineage and myeloid cell differentiation status in BM. **A.**Trajectory analysis of total BM cells. **B.** Lymphoid lineage cells reclustered and differential analysis using pseudotime. **C.** Myeloid subsetted and analyzed differentiation stage using sparse diffusion matrix. **D.** Sling-shot diffusion analysis identified clear dissociation of myeloid differentiation status. E. Density plot of cell-type specific markers of CMP (*Elane*), MM (*Ltf*) and BSN (*Tnfsf14*). Canonical cell type markers of CMP, MM and BSN were clearly expressed in each cell cluster location.

Neutrophils are the most abundant cells that play a central role in the innate immune response, and they are continually produced in the BM^21^. The transcription profile of the BM can reflect the myeloid differentiation and neutrophil maturation tendencies in the BM^22^. To reveal the direction of neutrophil differentiation in the scRNA dataset, we performed nonlinear dimensionality reduction of those data by restricting a sparse diffusion matrix of expression data to the eigenspace spanned by eigenvectors corresponding to the top diffusion matrix eigenvalues^23^. The diffusion map within the cell type showed a significant differentiation hierarchy and biased deposition of mature neutrophils **(Figure 2C)**. These tendencies were also visualized in Slingshot diffusion maps following diffusion components 1 and 2. The subclustering cells were arrayed in pseudotime **(Figure 2D)**.

This transcriptome velocity in myeloid differentiation was also confirmed by spatial deposition in a clustering map using a density plot **(Figure 2E)**. Representative markers of myeloblasts and promyelocytes, such as *Elane*, were clearly deposited in the CMP region **(Supplementary Figure 3A)**. Additionally, *Ltf*, *Cecam1* and *Olfm4*, well-known markers of MMs^22^, were exclusively expressed in MMs. Specifically, *Orm1*, which is not well known to be expressed in neutrophils, was observed to be specifically expressed in MMs, and *Camp*, which is known to be expressed in mature neutrophils, seemed to be expressed more specifically in MMs in our dataset **(Supplementary Figure 3B)**. The BSN marker genes *Adam8*, *Atp11a*, *Ctsd*, *Fcer1g*, *Lamp2*, *Mgam*, *Mmp8*, *Mmp9*, *Paur*, *Slc11a1*, *Timp2*, *Tnfaip6* and *Tnfsf14* (considered to be the most specific BSN marker) were dominantly expressed in the BSNs. In addition, *Il1a* and *Tgm2* showed specific expression in the BSN area **(Supplementary Figure 3C)**.

### Immature myeloid cells are more abundant in old BM than in younger BM

In the old BM, the hematopoietic microenvironment promotes myeloid expansion during physiological aging^24^. However, the proportion of committed myeloid cells in old BM is not fully understood. In this analysis, the proportion of committed precursors was significantly increased compared to that of mature cells in old BM, and this feature was most obvious among myeloid lineages.

Clear separation of specific cell populations between old and young mice was observed in 3D UMAP scattering **(Figure 3A)**. Regionally, a higher proportion of old BM clusters was observed in immature precursors in the neutrophil lineage, monocyte lineage and lymphoid lineage. More specifically, in the quantitative analysis, the proportion of MMs, iDCs and CLPs was increased, while the proportion of BSNs, MACs, MDCPs, pDCs and pre-B cells was decreased in old BM. Interestingly, there were no significant differences in young and middle-aged BM for most cell types and slight differences in iDCs, MDCPs and CMPs **(Figure 3B and Supplementary Figure 4)**. The significant changes in cell proportion between old and young mice indicated that these changes do not occur gradually with age but rather occur abruptly between middle and old age.

**Figure 3.**
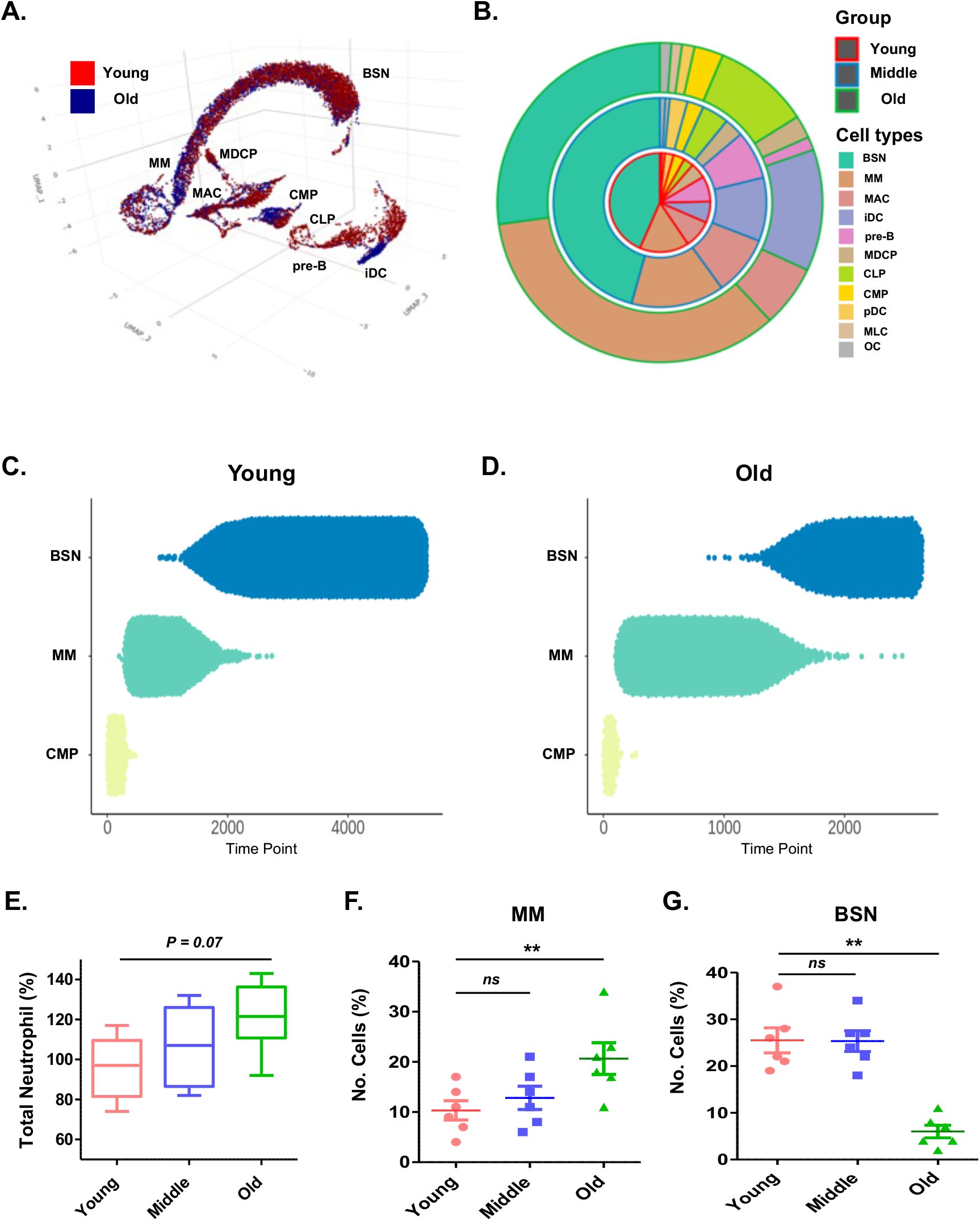
Immature myeloid cells are elevated in old BM. **A.** 3-dimensional (3D) UMAP analysis of young and old BM cells. Red dots indicate young cells, and blue dots indicate old cells. **B.** Circos plot describing age-dependent proportion changes in BM cells. Red, blue and green circles indicate young, middle and old BM, respectively. **C and D**. Diffusion analysis showed the myeloid differentiation status of young (C) and old BM (D). **E.** Flow cytometry analysis of neutrophils in the BM. **F and G**. Flow cytometry analysis of Ceacam1-positive MMs and Nlrp3-positive BSNs. (N=6 per group, *p value*: *ns* > 0.05, ** < 0.01 (one-way ANOVA))

Neutrophil lineages differentiate from hematopoietic stem cells (HSCs) and undergo differentiation to CMPs, MMs and BSNs in BM until they become functional neutrophils in the peripheral blood^25^. We observed that the increase in MMs and decrease in BSNs were the most significant changes in aging BM. We further analyzed the differentiation trajectory of neutrophils using diffusion analysis. In young BM, balanced neutrophil differentiation was observed, and the cell proportion was highest in BSNs **(Figure 3C)**, while the old BM appeared to be stagnant at the MM **(Figure 3D)**. This tendency was also observed by flow cytometry analysis. The total number of neutrophils was slightly increased in old BM **(Figure 3E)**. However, *Ceacam1*-positive MMs were highly increased in old BM **(Figure 3F)**, while *Nlrp3*-positive BSNs were significantly decreased in old BM **(Figure 3G)**.

These results suggested that a specific step of late-stage myeloid differentiation from MMs to BSNs was dysfunctional in old BM. Therefore, we further analyzed the cellular subtype of the neutrophil cluster using molecular subtyping.

### Molecular subtyping of neutrophils reveals alterations in secretory phenotypes with aging

We hypothesized age-dependent cell proportion changes in neutrophils due to changes in specific cell types rather than changes in whole cells. To identify specific cell types that have changed with aging, we reclustered the neutrophil lineages using dissociation signature genes.

In Figure 3B, the change of CMPs cell proportion with age was negligible, however, further analysis indicated that CMPs cluster can be internally divided into 7 subclusters with a significant dissociation signature genes and detected age-dependent proportional changes in the subclusters **(Figure 4A and B)**. In particular, the proportions of subclusters expressing *Gata2*+*ApoE*+ HSCs (cluster 0), CD63+*Ctsg*+ GMPs (cluster 1), *Irf8*+*Lgals1*+ CMPs (cluster 2) and *Mpo*+*Elane*+ GMPs (cluster 3) increased, while that of *Ermap*+*Add2*+ CMPs (Megakaryocyte–erythroid progenitor (MEP) committed cells, cluster 6) significantly decreased in the old BM compared to the young and middle groups **(Figure 4B and Supplementary Figure 5A and 6A)**. The functional enrichment of CMPs between young and old BM reflects the increase in Il-17 signaling/NK-cell differentiation and the decrease in cellular reprogramming-associated genes (Wnt) and inflammatory signaling **(Figure 4C)**.

**Figure 4.**
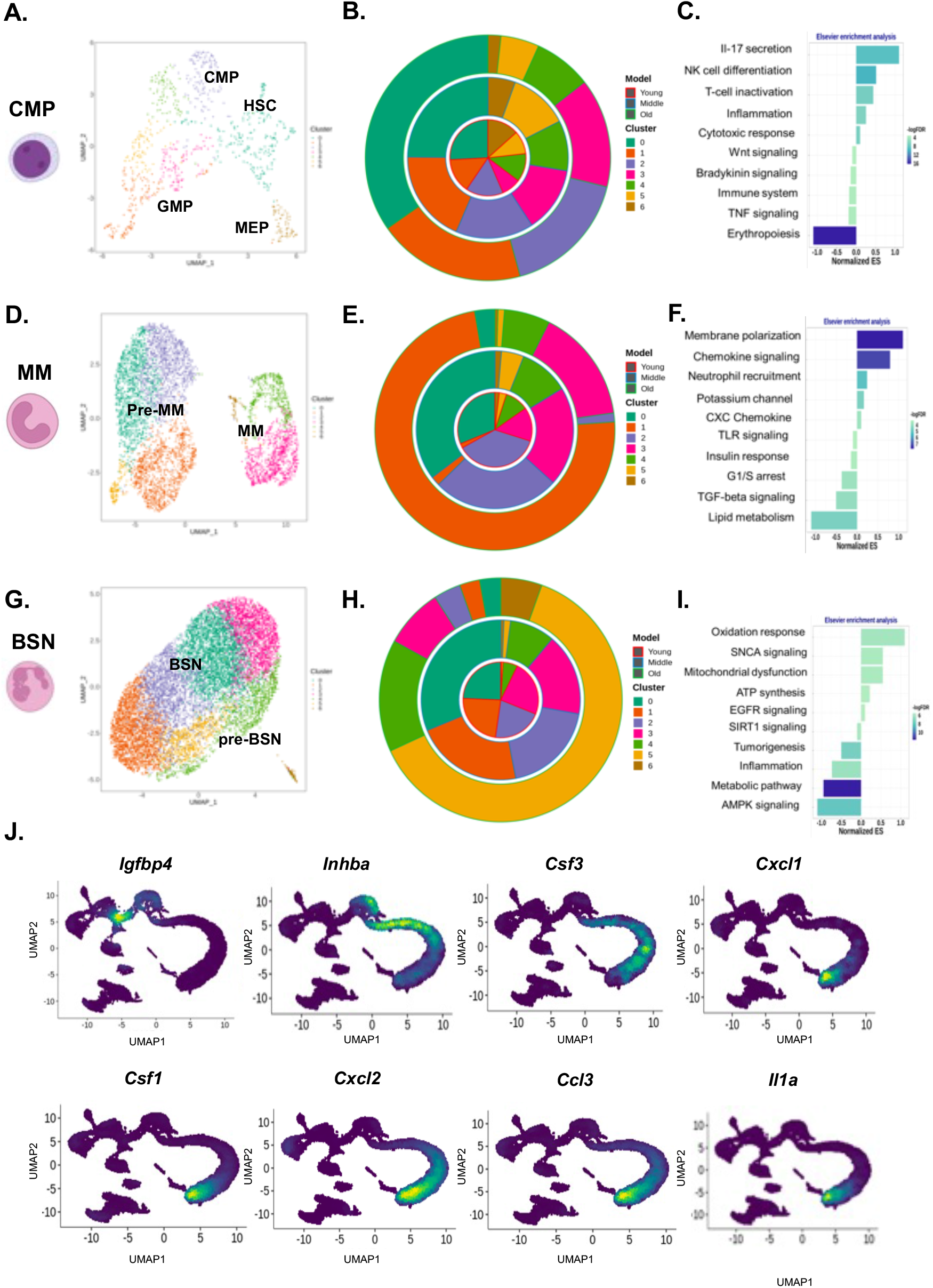
Unique subclusters of CMPs, MMs and BSNs were altered in old BM. **A-C.** UMAP, age-dependent cell proportion and enrichment analysis of total CMP subclustering. **D-F.** UMAP, age-dependent cell proportion and enrichment analysis of total MM subclustering. **D-F.** UMAP, age-dependent cell proportion and enrichment analysis of total BSN subclustering. **J**. Density plot visualizing differentially expressed secretory proteins specifically expressed in CMP (*Igfbp4*), MM (*Inhba*) and BSN (*Csf3, Cxcl1, Csf1, Cxcl2, Ccl3* and *Il1a*). Abbreviations: HSC (hematopoietic stem cell), MEP (megakaryocyte erythroid progenitor), pre-MM (MM precursor), pre-BSN (BSN precursor).

The increase of cell proportion in MMs in old BM was remarkable **(Figure 4B)**, and the composition of their subclusters were highly perturbed. MMs cluster can be subdivided into 7 clusters with clear dissociation signatures, which were divided into two particularly prominent groups **(Figure 4D)**. Committed myeloid progenitors robustly expressed mature neutrophil markers such as *Mmp8*, *Mmp9* and *Retnlg*. These cells were most common in young and middle BM (pre-MM, cluster 0, 24.7% and cluster 2, 22.2%) and were significantly decreased to 2.7% and 1.2% of cells, respectively, in old BM. In contrast, the proportion of cluster 1 that robustly expressed *Gadd45a* and *Cxcl2* prominently increased from 1.9% in young BM to 73.3% in old BM **(Figure 4E and Supplementary Figure 5B)**. Clusters 3, 4, and 6 (MM) showed a marked dissociation from the other clusters but did not show a notable cell proportion change with age. Cells of these clusters expressed genes related to neutrophil maturation (*Elane*, *Mpom*, *Vcam1*, etc.) and stem cell-like biological processes, such as cell cycle, development, or chromatin assembly regulatory factors (*Ube2c*, *Mki67*, *Tubb5*, *Pcna*, etc.) Indicating these clusters are the most committed but not highly differentiated. **(Supplementary Figure 6B)**. Corresponding with these results, enrichment analysis also showed a decrease in TGF-β signaling and Toll-like receptor (TLR) signaling and an increase in chemokine and neutrophil recruitment ligand signaling in old BM **(Figure 4F)**.

The proportion of BSNs decreased notably in old BM **(Figure 3B)**. BSNs cluster composed of comparably similar cells except for the clearly dissociated cluster 6 **(Figure 4G)**. The main factor of old BSN depletion is attributed to a decrease in the specific clusters. In the young and middle BSN, clusters 0 to 3 accounted for more than 88% of cells (young, 92.7%, middle 88.8%), whereas in the old BSN, this pattern was completely reversed, and clusters 0 to 3 accounted for less than 17% of cells **(Figure 4H and Supplementary Figure 5C)**.

Clusters 0 to 3 (marked BSN in Figure 4G), which were decreased in old BM, showed a highly functionalized neutrophil profile, expressing inflammatory response-associated factors (*Serpinb1a*, *Anxa1*, *Ccl6*, *Cybb*, *Chil3*, etc.) and interferon/interleukin signaling factors (*Plscr1*, *Ifitm6*, *Camp*, *Ltf*, *Ccl6*, etc.). Clusters 5 and 6 (marked pre-BSN in Figure 4G) robustly expressed activated neutrophil markers such as *Klf4*, *Jun*, *Fos* and *Tgfb1*. These clusters were a minor proportion in the young and middle BSN and became the main clusters in the old BSN **(Supplementary Figure 6C)**. These changes in the proportions of each cluster implied that there was a decrease in functional enrichment associated with neutrophil activation (AMPK signaling) and the inflammatory response in old BSNs **(Figure 4I)**.

Age-dependent changes in the cell proportions at each stage of neutrophil differentiation altered the neutrophil-derived secreted protein profile. Secreted interleukins (*Il-1a, Il-1b, Il-6, Il-7* and *Il-15*) were higher in young BM, while chemokines (*Ccl4, Ccl6, Ccl 27a, Cxcl1, Cxcl5*, etc.) were found to increase in old BM. Cytokines and growth factors were expressed differently in old BMs with increasing (*Areg, Vegfa, Hgf, Igfbp7*) and decreasing (*Egf, Igfbp4, Igfbp6*) patterns depending on the type **(Supplementary Figure 7)**. The age-dependent change in the expression of these secreted proteins was consistent with the changes in the proportions of CMP, MM, and BSN cells with age. As shown in the density plot, among the secreted proteins, *Igfbp4*, *Inhba* and *Csf3*, which were increased in old BM, were mainly expressed in CMP and MM, and *Cxcl1*, *Cxcl2*, *Ccl3*, *Il-1a*, and *Csf1*, which were decreased in old BM, were specifically expressed in highly differentiated BSNs **(Figure 4J)**.

Collectively, clustering and subclustering results indicated that myeloid lineage cells are accumulated in mitotic phase, and postmitotic phase cells are significantly decrease in old BM. Old BM could not provide circulating fully differentiated neutrophils, and as a result, the neutrophil-secreted protein profile appears to be altered in the old BM.

### Intercellular immune cell communication is altered in old BM immune cells

Alteration of intercellular communication by secretory and membrane-bound factors is critical for informing diverse cellular fate decisions, including decisions to activate the cell cycle, undergo apoptosis or differentiate along a specific lineage^26^. To identify the changes in intercellular communication that occur by aging, we used a curated comprehensive signaling molecule interaction database via CellChat^27^.

CellChat analysis revealed a marked difference in intercellular interactions between young and old BM. Overall, more complex and frequent interactions were observed in old BM than in young BM. Specifically, the major signals in young BM were derived from BSNs, MMs, MACs, CMPs and MDCPs, whereas in old BM, the signal density was increased and derived from a larger variety of cells, and autocrine signals were increased **(Figure 5A)**.

**Figure 5.**
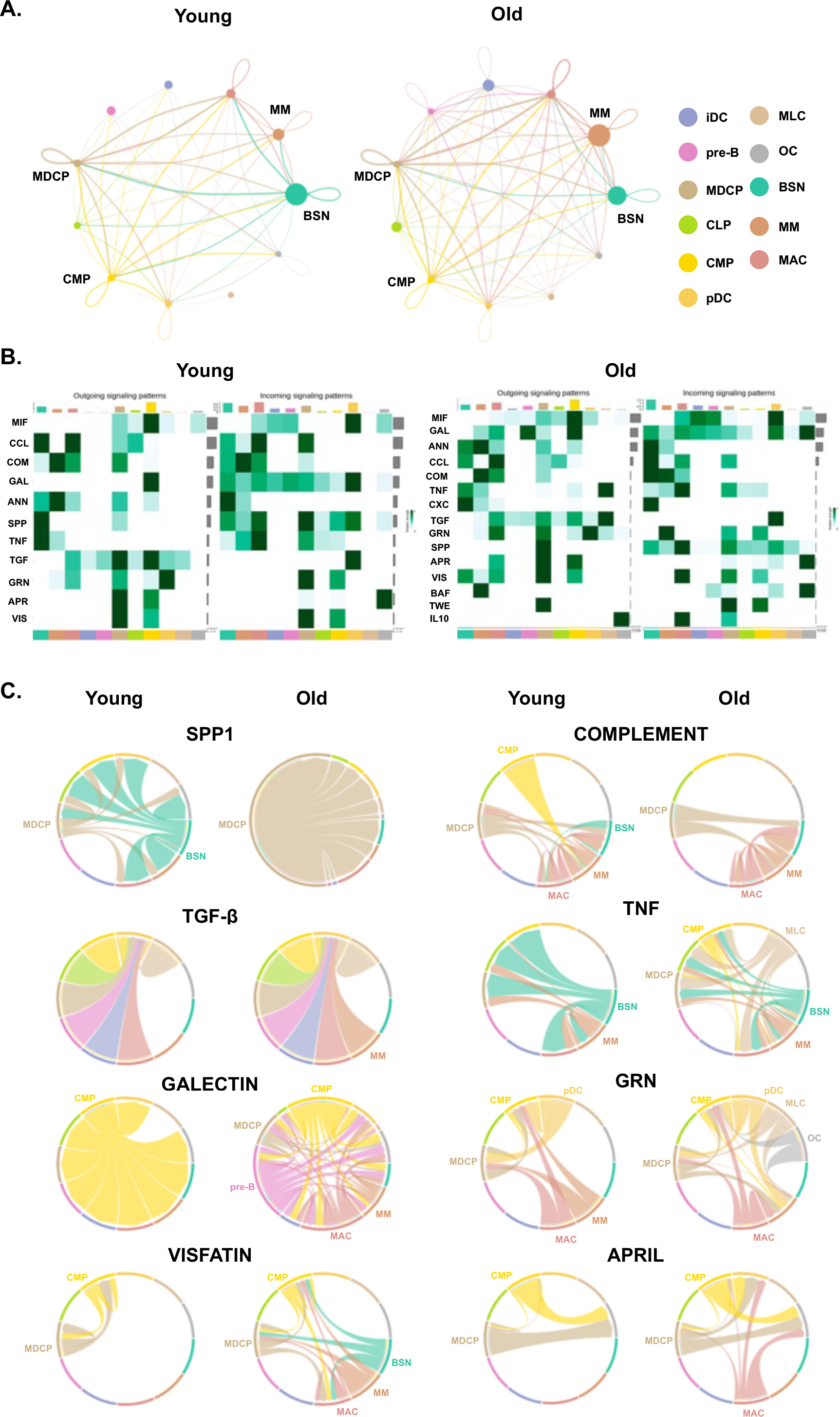
Intercellular immune cell communication was altered in old BM. **A.** Inferred total signaling networks of young and old BM. **B.** Heatmap depicting selected cell– cell interactions enriched in young and old BM. The color code indicates the predictive strength. **C.** Outgoing signaling patterns of inferred signaling networks between young and old BM. Color coding matched with cell type is consisted in all figures.

Among the 40 signaling pathway databases, significantly changed signaling pathways also showed significant differences between young and old BMs. Network centrality analysis of young BM identified 11 prominent signaling pathways, while that of old BM predicted 4 additional signaling pathways, CXCL, BAFF, TWEAK, and IL10, in addition to the 11 pathways of young BM. Even within the same signaling pathway, old BM showed more diversified signal-generating cells than young BM. For example, in young BM, TNF signaling is mainly emitted from BSNs and MMs, whereas in old BM, TNF signaling starts from 5 different cell types, BSNs, MMs, MDCPs, CMPs and MLCs. However, the receiving cells in the same signaling pathway did not changed significantly between the young and old groups **(Figure 5B)**.

Circos plot analysis of the outgoing signaling patterns of secreting cells predicted that SPP1 and COMPLEMNT signaling were repressed, while TNF, TGF-β, VISFATIN, GALECTIN, ARPIL (B lymphocyte stimulator (BLyS)-A PRoliferation-Inducing Ligand) and GRN (Granulin) signals were strengthened in old BM **(Figure 5C)**. In young BM, SPP1 was mainly secreted by BSNs and MDCPs, and SPP1 secretion was significantly simplified to MDCP alone in old BM. Additionally, the strong COMPLEMNT signals from CMPs and BSNs in young BM were extinguished in old BM. On the other hand, in old BM, the additional participation of MMs, BSNs and CMPs was remarkable in the generation of the TNF, TGF-β and VISFATIN signals. The GALECTIN and GRN signals in old BM showed a large change in the pre-B, MMs, and MACs.

Within the changes in signaling pathways, the ligand–receptor interactions in specific signaling pathways were also altered in old BM. Specifically, the ligand–receptor interactions of *Spp1*–*Cd44*, *Spp1–Itga5/Itgb1* and *Spp1–Itga4/Itgb1* in SPP1 signaling and *C3–Itgam/Itgb2* in COMPLEMENT signaling were predicted to be increased between young BSN and other cell types (BSN, MM and MAC). Molecular interactions in GALECTIN signaling were significantly increased between old MM and all cell types. Interestingly, in old BM, CXCL2 secreted from MM was predicted to interact with *Cxcr2* of BSNs, which has previously been reported as a neutrophil aging factor ^28,29^ **(Supplementary Figure 8)**.

These results imply that more idiotropic heterogeneous intercellular communications are developed in old BM, causing age-dependent immune-physiological changes such as skewing of the population balance of lymphoid and myeloid cells.

### Dysregulation of SPP1 signaling occurs in old BM due to a decrease in the BSN neutrophil population

The human immune cell population becomes increasingly biased toward the myeloid lineage in an age-related manner^30^. SPP1 coordinates the balance of the lymphoid and myeloid populations in BM^31^ and is also known to regulate HSC pool size, stem cell homing, trans-marrow migration and engraftment^32,33^. Previously, depletion of SPP1 was mainly caused by a decrease in secretion by osteogenic cells in BM^34^, and depletion of SPP1 in old BM induced senescence phenotypes in HSCs, including loss of polarity and myeloid skewing^35^.

The expression level of *Spp1* significantly decreases by aging in both humans and mice. The *Spp1* expression level in total BM was decreased significantly in the old BM compared with young and middle BM **(Figure 6A)**, which was consistently reproduced in bulk-RNA sequencing data of human and mouse BM **(Figure 6B, C)**.

**Figure 6.**
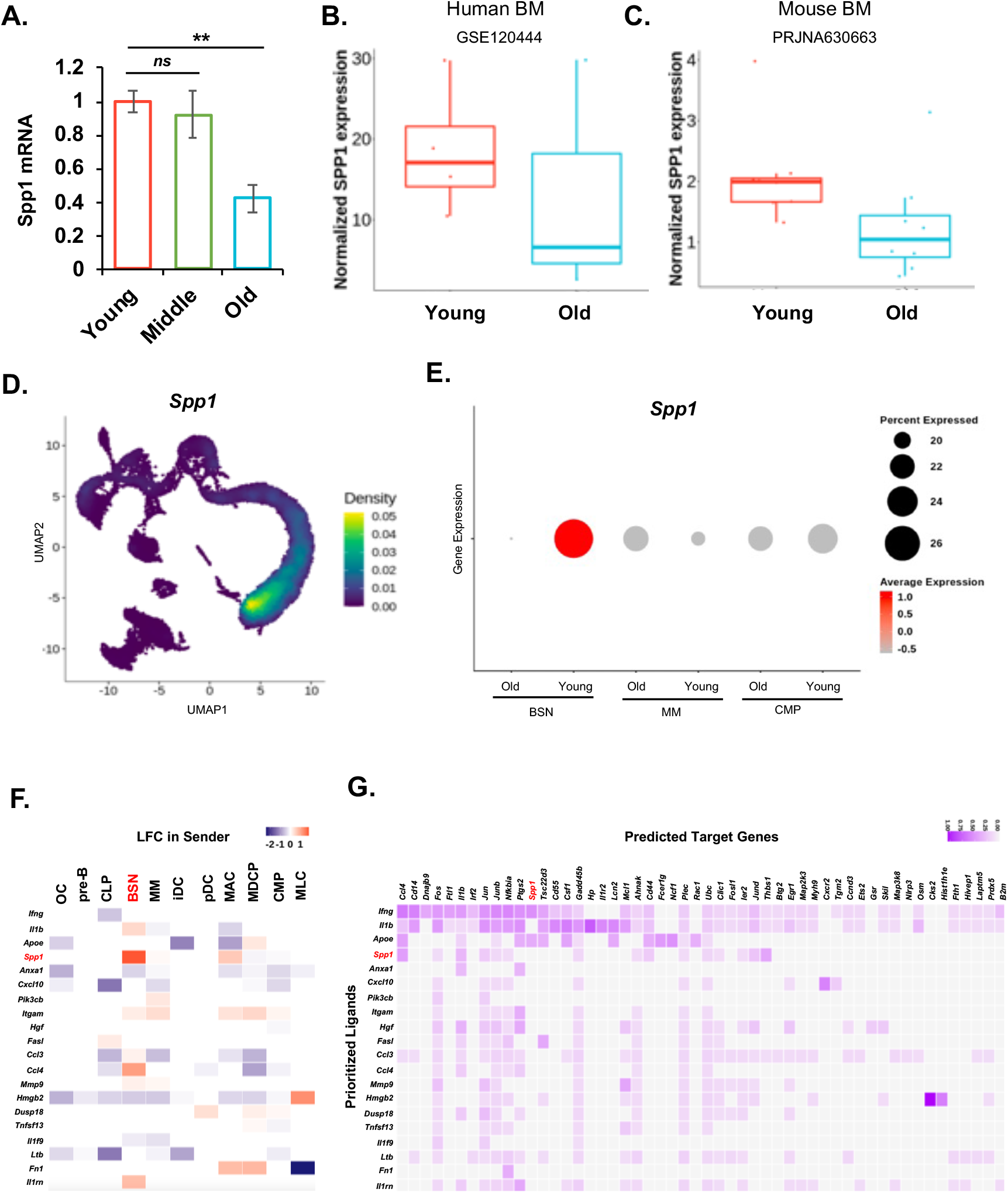
Depletion of SPP1 in old BM caused by depletion of BSNs. **A.** Real-time qPCR analysis of Spp1 in young, middle-aged and old BM. (N=3 per group, *p value*: *ns* > 0.05, ** < 0.01 (one-way ANOVA)) **B and C.** Public bulk RNA-seq database analysis of *Spp1* expression in humans (B) and mice (C). **D.** Density plot visualizing Spp1 expression in total BM. **E.** Bubble chart showing the age-dependent expression level and cell proportion of Spp1 in CMP, MM and BSN. **F.** Inferred PPI networks between young and old BM. Increase in LFC denoting the predictive activity in young BM. **G.** Ligand–receptor matrix denoting the regulatory potential between old and young BM.

*Spp1* was exclusively expressed in BSNs in the total immune cell population **(Figure 6D)**. The major cell population (over 26%) of young BSNs highly expressed *Spp1* but in old BSNs its expression was significantly diminished **(Figure 6E)**.

To define the influence of the depletion of SPP1-generating cells in old BM, we used a ligand–receptor interactome-based protein–protein interaction (PPI) predictive model called NichNet^36^. In old BM, *Ifng*, *Il1b*, *Apoe* and *Spp1* levels were lower than those in young BM **(Figure 6F and Supplementary Figure 9A)**. Notably, *Spp1* was the highest priority ligand in BSNs and was predicted to target *Ccl4*, *Fos*, *Il1b*, *Jun*, *Junb*, *Nfkbia*, *Gadd45b*, *Ahnak*, *Cd44*, *Plec*, *Ubc*, *Clic1*, *ler2*, *Jund* and *Thbs1*, which were enriched in the regulation of the cell cycle and inflammatory response **(Figure 6G and Supplementary Figure 9B)**. Predictive PPI visualized interactions between SPP1 and integrin family members (such as *Itga4*, *Itgb3*, *Itga8*, etc.) in BSNs and ANXA1 signaling in MMs **(Supplementary Figure 9C)**. Interestingly, *Spp1* was predicted to be regulated by *Ifng* and Apoe in young BM, and these mechanisms might be downregulated in old BM **(Figure 6G)**.

Collectively, depletion of SPP1 signaling in old BM is linked to a decrease in BSN and an increase in MM populations in an age-associated manner. Additionally, the specific alteration of the cytokine-producing neutrophil population results in the perturbation of normal signaling pathways, which are key regulatory factors for the balance of hematopoiesis.

## Discussion

In this study, we performed scRNA-seq analysis of total BM immune cells except for nonimmune cells such as adipocytes and RBCs that were excluded by density gradient centrifugation or hypotonic shock. There have been many reports of bulk RNA-seq analysis or scRNA-seq of FACS-sorted specific groups of BM cells. Unfortunately, however, there have been no deep analyses of scRNA-seq of total BM cells representing age-dependent changes in the total BM immune cell population profile.

With this approach, we found that young BM showed a well-organized differentiation status from the precursor to mature immune cells in which myeloid lineage cells were predominant, which is coincident with previous reports^37,38^. We identified a high proportion of committed but functionally immature myeloid lineage cells in old BM, including MMs in the neutrophil lineage and iDCs in the monocyte lineage. Subclustering of neutrophils to identify aging-sensitive cell types revealed that the increase in MMs and decrease in BSNs in old BM was caused by fluctuations in the proportion of specific immature cell types rather than all cells. These proportion changes revealed perturbation of intercellular communications in the immune cells in old BM. Specifically, many signals derived from BSNs in young BM disappeared or were substituted by other cells, including MMs. Collectively, our data strongly suggest that gradual deposition of immature neutrophils is a core feature of old BM and a potential contributor to dysfunction of the immune response in elderly individuals.

Aging-associated immune cell dysfunction of BM is frequently explained by the alteration of early fate determination in HSCs and CMP/CLPs^39–41^. With advances in single-cell transcriptome analysis, previous studies analyzed age-associated changes in BM using FACS-sorted HSCs^42,43^, although the proportion of high-order HSCs in BM is less than 0.01%. However, few studies have focused on the mature myeloid lineage in aging, even with high abundance^44^. Recently, a bulk omics profiling study of BM neutrophils identified aging- and sex-dependent functional differences caused by epigenomic alterations^45^. However, bulk omics analysis has limitations in analyzing cell proportion changes in mature myeloid lineages and associated functional changes by aging. Our data provide the first representation of single-cell-level age-dependent immune cell profiling from freshly isolated BM. Total single-cell clusters can be defined as one of 11 cell types and showed a high abundance of the myeloid lineage (over 87% of cells). Relatively few lymphoid cells were detected, at approximately 11.4% of the cells, while pre-B cells represented 6.7% of the cells. Subclustering analysis of the lymphoid lineage showed that the T-cell population is incorporated into CLPs. In myeloid cells, neutrophils showed the highest population, followed by MACs and DCs. This single-cell profile showed a similar cell composition as that previously reported in young BM^37^.

The prediction of the differentiation direction of the myeloid lineage in our dataset using trajectory analysis well reflected a previously known differentiation tendency and the newly suggested cellular proportion of each differentiation stage. Neutrophil maturation consists of a mitotic phase (CMP to MM) and a postmitotic phase (MM to BSN)^46^. Our data suggested that differentiation failure of old neutrophils occurs at the postmitotic phase, leading to the accumulation of immature dysfunctional cells in aged BM. These results indicated that dysfunction of the innate immune reaction in elderly individuals may be due to accumulation of the immature myeloid population and depletion of functional immune cells. Several studies in humans have shown that the number of neutrophils in the blood is slightly lowered, but neutrophil function is notably decreased in healthy elderly individuals^47,48^.

3D-UMAP analysis revealed prominent proportional changes in the specific subpopulations within each cluster with aging. Subclustering of each cell type effectively characterized these specific subpopulations, and these cells showed significant intercluster differences in gene expression profiles. The most significantly changed subcluster in MMs expressed the DNA damage-inducing protein *Gadd45a* along with neutrophil maturation markers such as *Cxcl2* and MMPs. Considering the increase in the proportion of this subcluster from 1.2% in young BM to 73% in old BM, it is thought that the differentiation to BSNs, a postmitotic cluster, is not progressing due to the increase in the proportion of DNA damaged MMs cluster cells. The most abundant BSNs in young BM expressed *Serpinb1a*, previously reported to prevent cathepsin G-dependent neutrophil cell death^49^, and *Anxa1*, which is known as a regulator of neutrophil maturation and migration^50^. Collectively, these results suggest that the population of mature BSNs is reduced and that the rest of the neutrophils are mostly tied to undifferentiated MMs clusters.

Alteration of the cell proportion directly influenced the intercellular communication landscape of BM. We used two different cell–cell communication analysis databases, CellChat and NicheNet. Using these analyses, we newly suggested that the decrease in the proportion of BSNs could be the main contributing factor to SPP1 depletion in old BM. Aging altered the cellular composition of the HSC niche, depletion of SPP1 signaling skewed myeloid imbalance during pathogenic conditions^31^, and treatment with SPP1 led to phenotypical and functional rejuvenation^35^. Despite the broad role of SPP1 signaling in hematopoiesis, only osteogenic lineage cells have received attention as the major source of SPP1 in BM^51^. In our analysis, *Spp1* was specifically expressed in BSNs in young BM, and the aging-associated decrease in the proportion of BSNs is closely related to a decrease in SPP1 concentration in the BM. Moreover, the weak expression of *Spp1* even for BSNs present in old BM indicates that the characteristics of old BSNs are different from those of young BSNs.

Previous studies reported a direct correlation between IFN-γ and SPP1 levels in blood from human patients with mycobacterial infection or schizophrenia^52,53^. In our analysis, *Ifng* and *Apoe* were predicted to be SPP1 signaling inducers in young BM. Although a direct correlation between *Apoe* and *Spp1* has not yet been reported, in large-scale transcriptome analysis studies, *Spp1*, *Apoe* and *Ifng* showed high coexpression rates, especially in Alzheimer’s disease-involved microglial cells^54^. Collectively, the correlation between systemic depletion of SPP1 levels in elderly individuals and geriatric neurological pathologies may be linked with the depletion of BSN in old BM.

These predictive alterations of intercellular communication are strongly associated with the first line of viral immunity. An uncontrolled inflammatory response to COVID-19 is a critical factor for severe patients, especially in elderly individuals^55^. The majority of patients with severe COVID-19 are reported to have substantially elevated serum levels of proinflammatory cytokines and chemokines, including CCL3 and TNF^56^. In elderly patients, reactive oxygen species (ROS) levels increase with age and can potentiate NLRP3 (classified as a marker of MMs in this dataset) activation-associated pyroptosis, and the release of DAMPs further heightens inflammation and COVID-19 pathogenesis^57,58^. Additionally, depletion of pDCs may cause reduced TLR7 expression and diminished viral immunity in elderly individuals^59^. Collectively, increase in MMs in old BM may be linked with the higher incidence and severity of COVID-19 in older patients.

In summary, we analyzed young, middle-aged and old BM using scRNA-seq to reveal alterations in the immune cell proportion landscape associated with aging. This dataset highlights the overall myeloid skewing and repression of the functional commitment of neutrophils in old BM. Indeed, alteration of the neutrophil population with aging appears likely to be one mechanism that causes immunomodulatory defects in elderly individuals. These results may provide a new connection between aging-associated innate immune dysfunction and the systemic association between BM and other age-sensitive organs, such as the neuronal system.

## Materials and Methods

### In vivo experimental details

The mice used for all experiments were C57BL/6J wild-type mice acquired from KRIBB (Korea Institute of Bioscience and Biotechnology, Korea), where the animals were housed in the same housing facility from birth until purchase. Then, the animals were acclimatized together at the animal housing facility of Seoul National University for at least 2 weeks before experimental procedures. Mice were kept in a 12-hour light–dark cycle, with an ambient temperature range within 20–24 °C; humidity ranged from 30% to 70%. All male mice from the ‘young’ group were aged approximately 3 months, those from the ‘middle’ group were aged 12 months, and those from the ‘aged’ group were aged 20 months. All animal experiments performed in this study were approved by the Animal Care and Use Committee of Seoul National University under animal protocol number SNU-210202-5-3.

To prepare fresh bone marrow cells, we isolated total bone marrow cells as reported previously^60^. Briefly, bone marrow cells were extracted from mouse tibias and femurs, and immune cells were isolated using RBC lysis buffer (Miltenyi Biotec, no. 130-094-183) and Ficoll centrifugation, albeit with no vortexing step to avoid unscheduled immune cell activation. The harvested cells were washed several times with washing buffer to remove dead cells to achieve over 90% live-cell proportion. All harvested cells were immediately ready for scRNA-seq preparation or stored in deep freezers for subsequent experiments.

### Single-cell experiment and data extraction

Single cells were encapsulated into emulsion droplets using a Chromium Controller (10x Genomics). Single cells, reagents and a single gel bead containing barcoded oligonucleotides are encapsulated into nanoliter-scale GEMs (gel bead in emulsion) using the Next GEM Technology. The library kit used Chromium Next GEM Single Cell 3p RNA library v3.1. Single Cell 3’ GEX and feature barcode libraries were sequenced on the Illumina sequencing system for paired-end sequencing. Cells were then loaded in each sample with a target output of more than 4,000 cells. scRNA-Seq data were demultiplexed, aligned to the mouse genome, version mm10, and UMI-collapsed with the CellRanger toolkit version 6.0.1 available from 10x Genomics with default parameters. The ‘mkfastq’ module in CellRanger demultiplexes raw base call (BCL) files generated by Illumina sequencers into FASTQ files. The total number of bases, reads, GC (%), Q20 (%), and Q30 (%) were calculated for three samples (young, 3 months; middle, 12 months; old, 20 months). The total produced reads were 533,762,334, 305,564,464 and 332,155,986 in young, middle and old mouse models, respectively. The mean Q30 ratios were 91.99%, 93.86% and 93.99% in the young, middle and old mouse models, respectively.

### Data preprocessing and doublet elimination

Gene expression was processed using the “Seurat, v.4.1.1.” R package^61^. We removed genes that were not expressed in at least 10 cells and then cells that did not have at least 200 detected genes. For feature filtering, we removed cells with fewer than 2,500 counts and fewer than 5% mitochondria. Gene expression was represented as the fraction of its UMI count with respect to total UMI in the cell and normalized using size factor normalization such that every cell has 10,000 counts (TP10K) and log transformed. The computationally predicted doublet cells were eliminated by the “DoubletFinder, v2.0.” R package^62^.

### Dimensional Reduction and Clustering

We performed dimensionality reduction using gene expression data for a subset of 2000 highly variable genes. The variable genes were selected based on the dispersion of binned variance to mean expression ratios using the FindVariableGenes function of the Seurat package followed by filtering of cell-cycle, ribosomal protein, and mitochondrial genes. Next, we performed principal component analysis (PCA) and reduced the data to the top 50 PCA components (number of components was chosen based on standard deviations of the principal components – in a plateau region of an “elbow plot”). We used graph-based clustering of the PCA reduced data with the Louvain method after computing a shared nearest neighbor graph. We visualized the clusters on a 2D map produced with t-distributed stochastic neighbor embedding (t-SNE) and uniform manifold approximation and projection (UMAP). For reclustering, we applied the same procedure of differentially expressed genes in the analyzed cluster, dimensionality reduction, and clustering to the restricted set of data (usually restricted to one initial cluster). Three-dimensional plots of young and old were plotted using the plotly R package.

### Dissociation and molecular subtyping analysis

We checked the possible impact of dissociation by several analyses. First, we reran the clustering of our dataset but after removing the known dissociation artifact genes from the data matrix^63^. We did not observe substantial clustering changes in this approach. Specifically, the number of clusters remained the same, and 98% of cell pairs had the same cluster assignment. Second, we performed binning of cells into “clusters” based on the expression of only dissociation signature scores (with the same bin sizes as the real clusters): the adjusted random index between our full clustering and this binning was 0.009 (for equivalent clustering, the value is 1). This indicated that our global clustering was not driven by dissociation signature expression. Third, we repeated this test for clusters that were further subclustered. This indicated that only the reclustering of progenitor and differentiated cells was possibly partially correlated with the dissociation signature or other biological processes with correlated signatures (we note that immediate early gene expression is the main feature of the “dissociation signature” and can also be a biological phenomenon).

### Cell type annotation and composition

Cell types with functional identities were assigned to cell clusters based on differential expression signatures derived from statistics of the log likelihood ratio test. To define the cell clusters, we performed principal component analysis on the most variable genes between cells, followed by Louvain and Leiden graph-based clustering. We extracted significantly differentially expressed genes in each of the 15 original clusters from the bone marrow samples. Next, we computed how many cell types mapped to each individual cluster. For elaborate mapping between clusters and cell type, we used three algorithms with the well-known databases CellMarker, Mouse Cell Atlas (MCA), and Single R. Finally, we confirmed clusters and important markers using manual curation based on the consensus of three experts. When assigning the subcluster identity, we further scored against signature genes of subclusters that consisted of up to most differentially expressed subcluster genes.

### Differential expression analysis (DEA)

To identify remarkable markers in each cluster, we applied the Wilcoxon Rank-Sum Test (using FindMarkers of Seurat package) to find genes that had significantly different RNA-seq TP10K expression compared to the remaining clusters. The selection criteria were 0.05 levels (FDR-q after multiple hypothesis testing correction) of statistical significance and more than 2-fold change between each cluster. We performed differential gene expression analysis on each cell type in young, middle-aged and old mice. We used a generalized linear model treating the three aging groups (young, middle and old) as a categorical variable by the “DESeq2” R package. We applied a false-discovery rate (FDR) threshold of 0.01 and a 4-fold change between aging groups.

### Density of differential expression across multiple cell types

Due to the sparsity observed in single-cell data, the visualization of expression among the cell types is frequently affected and unclear, especially basic dimension plot when it is overlaid with clustering to annotate cell types. We visualized data from the relative gene expression profiles of single cells using the “Nebulosa” R package^64^. It recovers the signal from dropped-out features by incorporating the similarity between cells, allowing a “convolution” of the cell features using kernel density estimation.

### Overrepresentation analysis for gene set enrichment

Overrepresentation analysis was performed to identify biological pathways that were differentially expressed. The database for annotation used Elsevier pathway and Gene Ontology terms to facilitate high-throughput gene functional analysis. The overrepresentation analysis enriches the biological information of individual genes and can be used to annotate gene-related biological mechanisms using standardized gene terminology. The statistical significance was determined using a hypergeometric test. For multiple hypothesis correction, a statistical test was performed using Benjamin-Hochberg’s method.

### Trajectory and diffusion map analysis

To define the transcriptional dynamics of a temporal process of cell differentiation, we used an unsupervised algorithm that increases the temporal resolution of transcriptome dynamics using single-cell RNA-Seq data collected at multiple time points using the “monocle3” R package^20^. Applying the methodology to total bone marrow cells and myeloid progenitor-derived cells, we identified kinetic time (pseudotime) lineage changes in the expression of previously known cell differentiation markers. We performed nonlinear dimensionality reduction of scRNA-seq data by restricting a sparse diffusion matrix of expression data to the eigenspace spanned by eigenvectors corresponding to the top diffusion matrix eigenvalues. (Rare cell proportions with fewer than 1% OC and MLCs were excluded from the diffusion map analysis.)

We used the “destiny” R package, where we used the local estimation of Gaussian kernel width, and the number of nearest neighbors for diffusion matrix approximation was set to the smaller value between the square root of the number of all single cells in the data and 100^23^. The diffusion matrix was constructed on sets of variable genes computed with the same procedure as the one used for clustering and subclustering of the data, such that variable genes were recomputed for each diffusion map. We visualized diffusion maps following diffusion components 1 and 2. The subclustering cells were arrayed at the pseudotime point.

### Cell-to-cell communication and network inference analysis

We evaluated cellular communication across various cell types through quantitative inference and network analysis of intercellular communication from single-cell RNA-sequencing data. We performed analysis and visualized data from single cells using the “CellChat” R package. We calculated the model probability of cell–cell communication by integrating gene expression with prior knowledge of a total of 40 remarkable signaling pathways of the interactions between ligands, receptors and their cofactors as senders, mediators and receivers of signaling architectures. We performed network inference analysis using the “Nichenetr” R package. We calculated strong ligand activity, predictive target genes, and the network of interacting genes in signaling pathways.

### External validation of SPP1 expression in a public dataset

We conducted verification with external data to examine whether the expression level of the SPP1 gene differs between young and old individuals. The two datasets PRJNA630663 and GSE120444 in mice and humans, respectively. The data of PRJNA630663 were obtained by extracting neutrophils from the bone marrow of eight young (4 months) and eight old (20 months) mice. The GSE120444 data were obtained by extracting mononuclear cells from bone marrow aspirates from four young (age < 47) and three old (age > 50) humans. The statistical significance of the difference in SPP1 expression was calculated with the Wilcoxon rank sum test for nonparametric distribution using the basic function of the R program.

### Flow cytometry and Gene expression

Neutrophils were isolated from other bone-marrow cells using MACS (Miltenyi Biotec, no. 130-097-658). Viability and yield were assessed using trypan blue exclusion and an automated COUNTESS cell counter (Thermo Fisher Scientific). MACS-purified neutrophils were then stained using anti-Ly-6G-PE (Invitrogen, no. 12-9668-82) and CD11b-FITC (Invitrogen, no. 11-0112-82) at a 1:50 dilution according to the manufacturer’s instructions. Stained cells were then analyzed by flow cytometry (FACSVerse, BD bioscience).

Total RNA was extracted from cultured cells using an RNeasy Mini Kit (#74104, Qiagen) and reverse transcribed into complementary DNA using a PrimeScript™ RT Master Mix (RR036A, TaKaRa Bio) in accordance with the manufacturer’s protocol. TB-Green® Premix Ex Taq™ (RR420A, TaKaRa Bio) was used for quantitative PCR (qPCR) in an Applied Biosystems™ 7500 RT-PCR system. Results were normalized to Gapdh expression.

### Data availability

The sequencing data of the main analysis have been submitted to the Sequence Read Archive (SRA) accessible through BioProject PRJNA868133. The supported data analyzed in this manuscript are publicly available from the GEO (www.ncbi.nlm.nih.gov/geo/) under accession number GSE120444 (human bulk RNA-seq data) and SRA (https://www.ncbi.nlm.nih.gov/sra) under accession number PRJNA630663 (mouse bulk RNA-seq data).

### Code availability

The main analysis for processing data and visualization of single-cell RNA-seq are available from Seurat (https://rdrr.io/cran/Seurat/man/). Trajectory analysis is available from monocle3 (https://cole-trapnell-lab.github.io/monocle3/). Cell–cell communication analysis is available from GitHub source code repositories with cellchat (https://github.com/sqjin/CellChat) and nichenet (https://github.com/saeyslab/nichenetr).

## Supplementary Figures

**Supplementary Figure 1.**
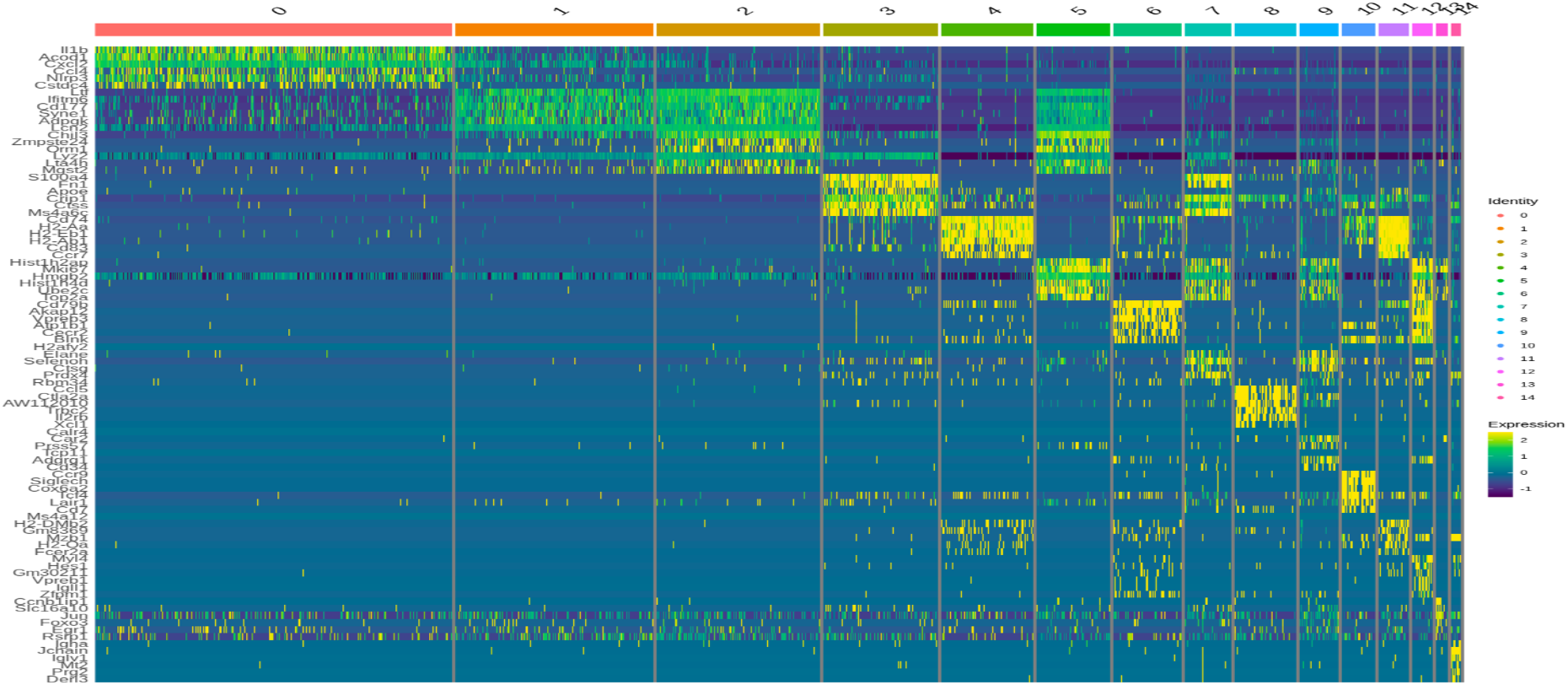
Heatmap of cell-type-specific gene signatures of total BM cells.

**Supplementary Figure 2.**
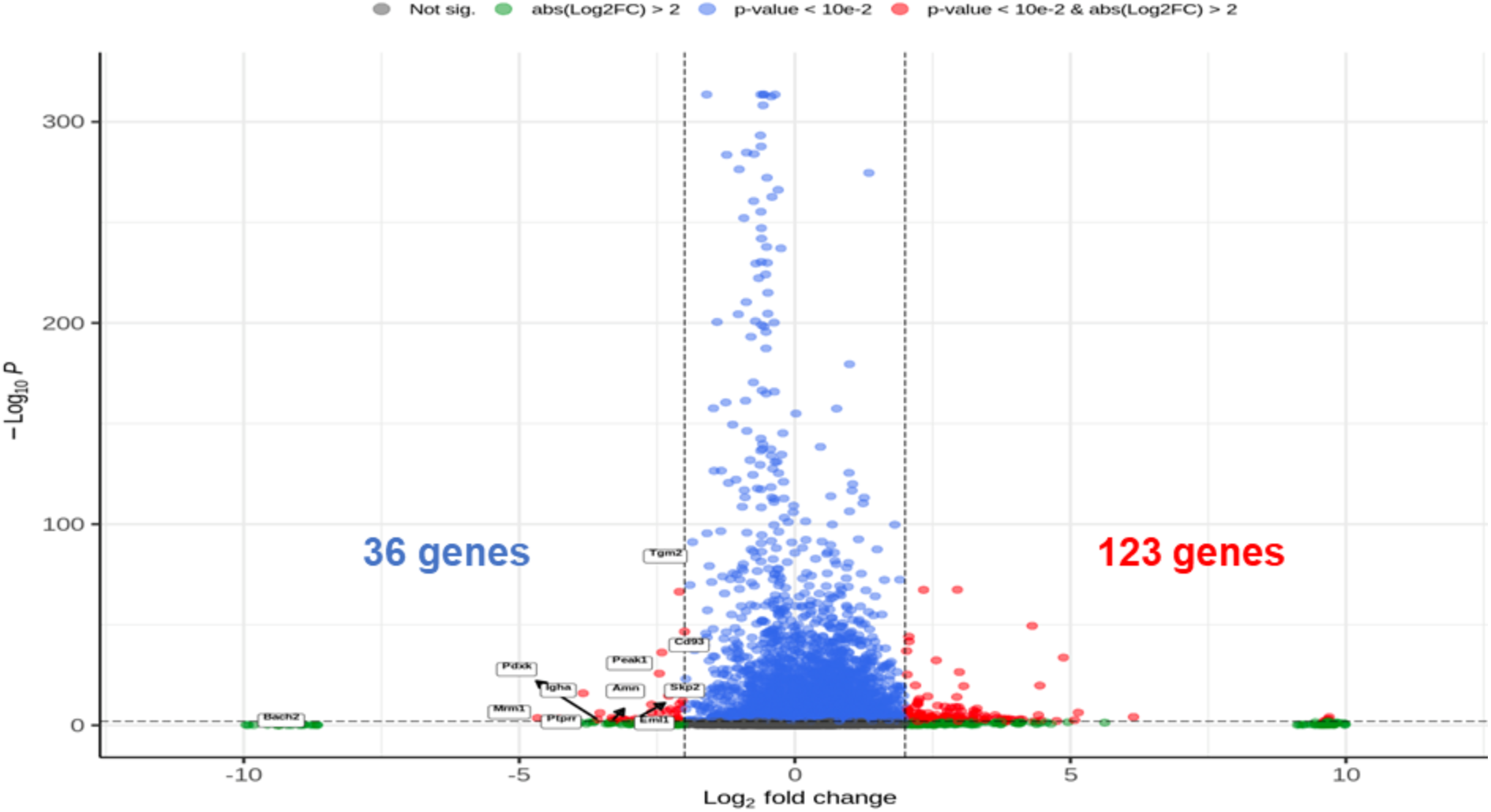
Differential gene expression profile of old versus young BM.

**Supplementary Figure 3.**
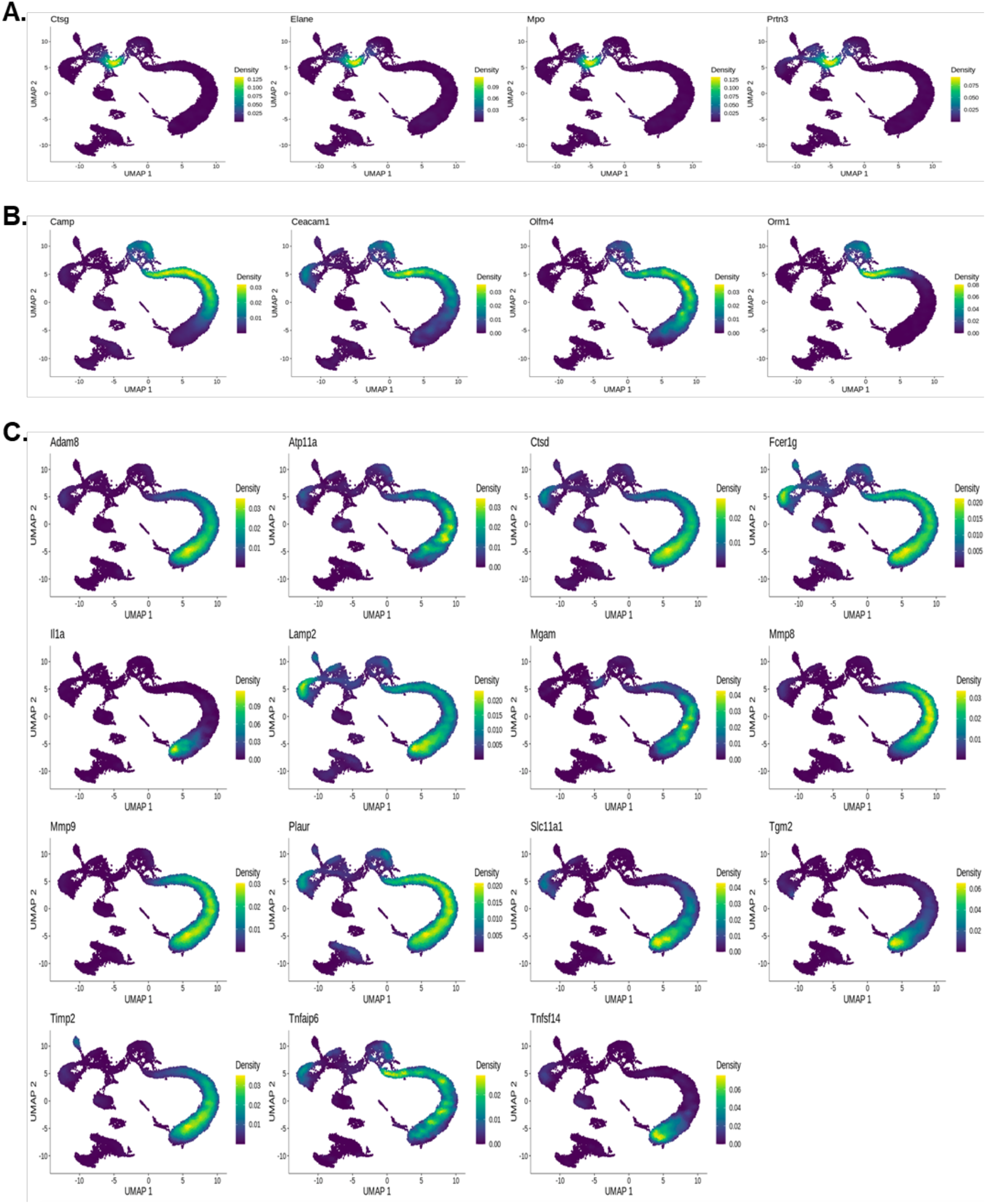
Density plot of CMP-, MM- and BSN-specific markers.

**Supplementary Figure 4.**
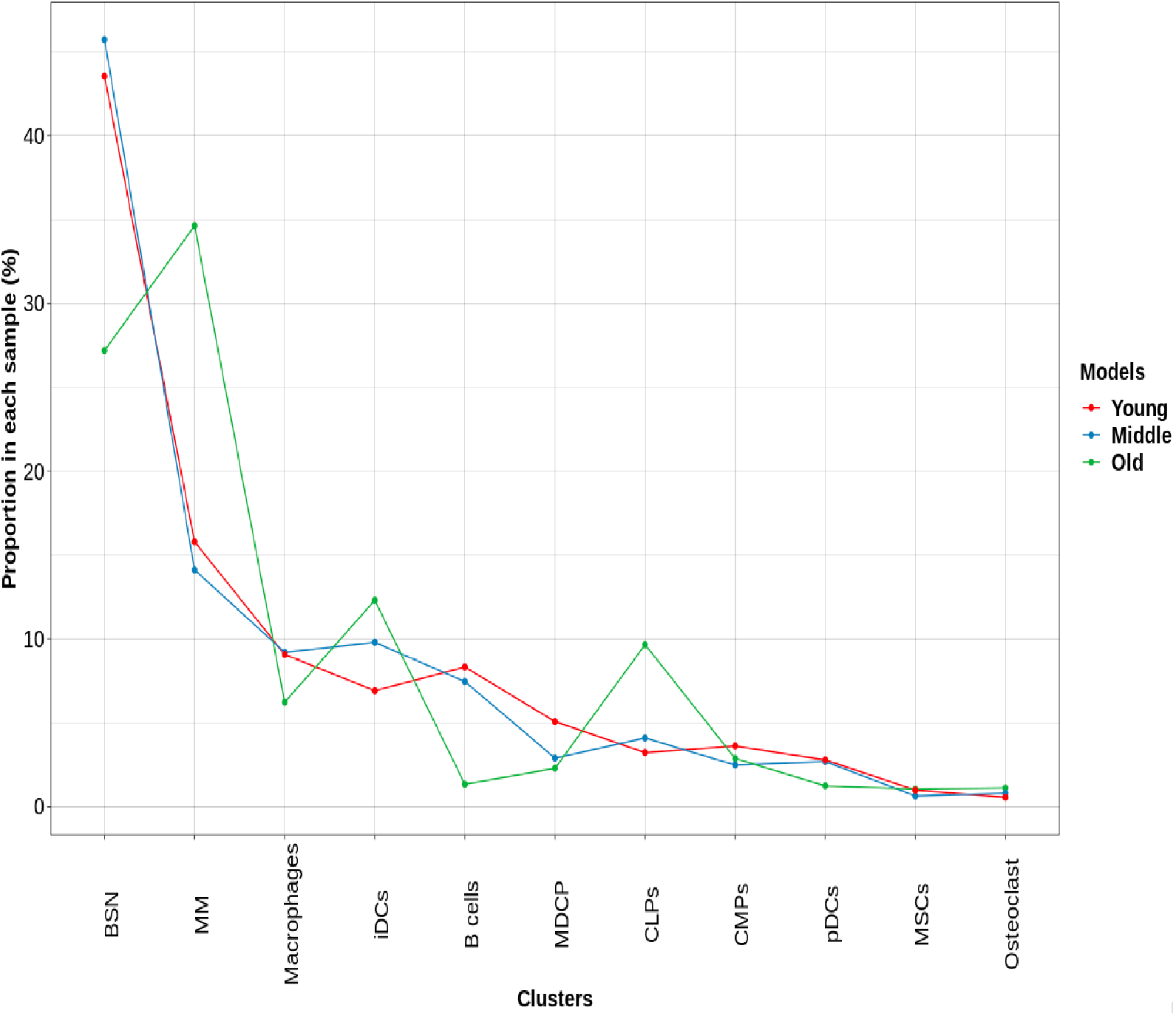
Age-associated cell-type population changes.

**Supplementary Figure 5.**
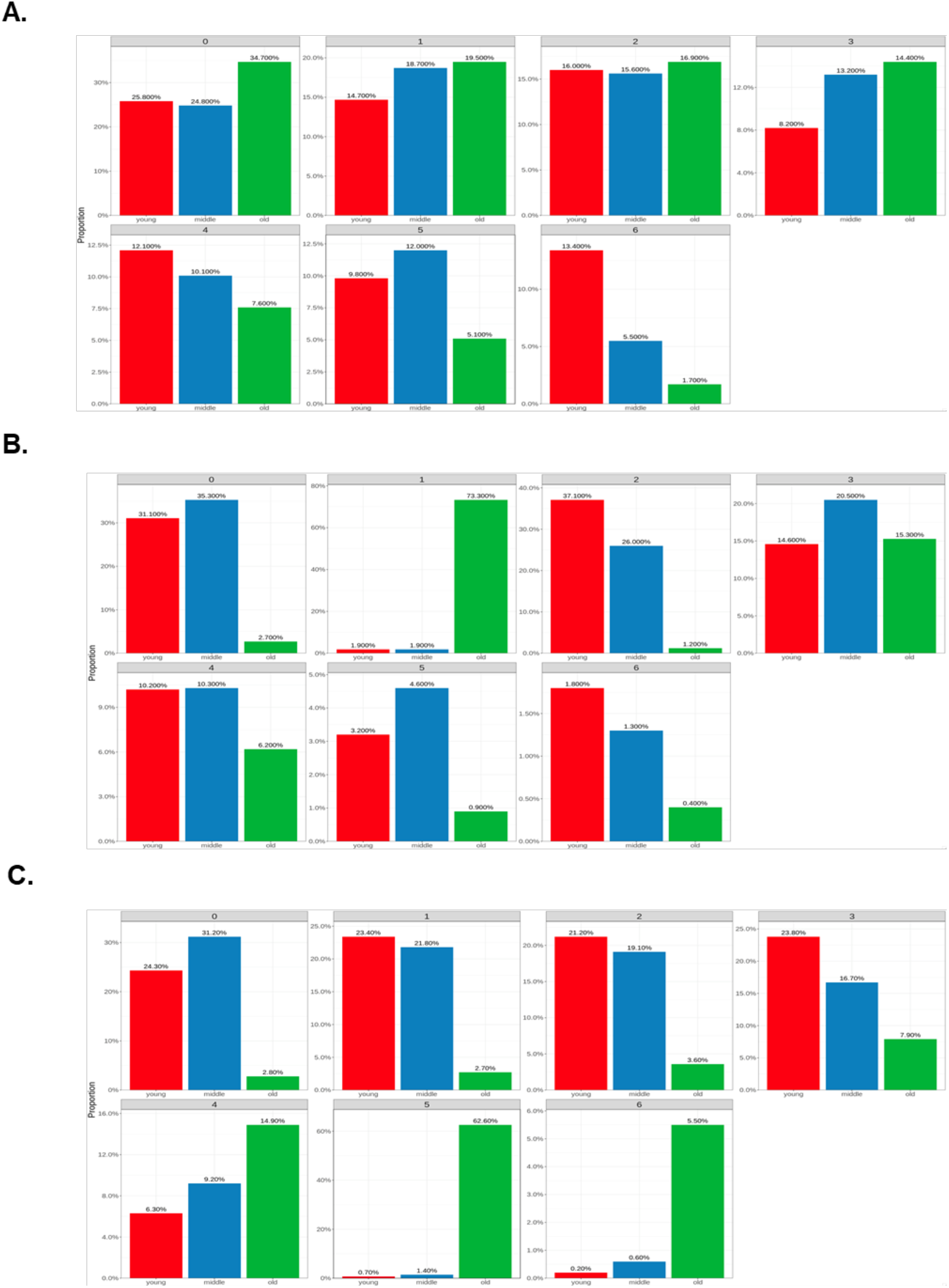
Age-associated population changes of CMP, MM and BSN subclusters. Red bar indicate young, blue bar means middle and green bar indicate old group.

**Supplementary Figure 6.**
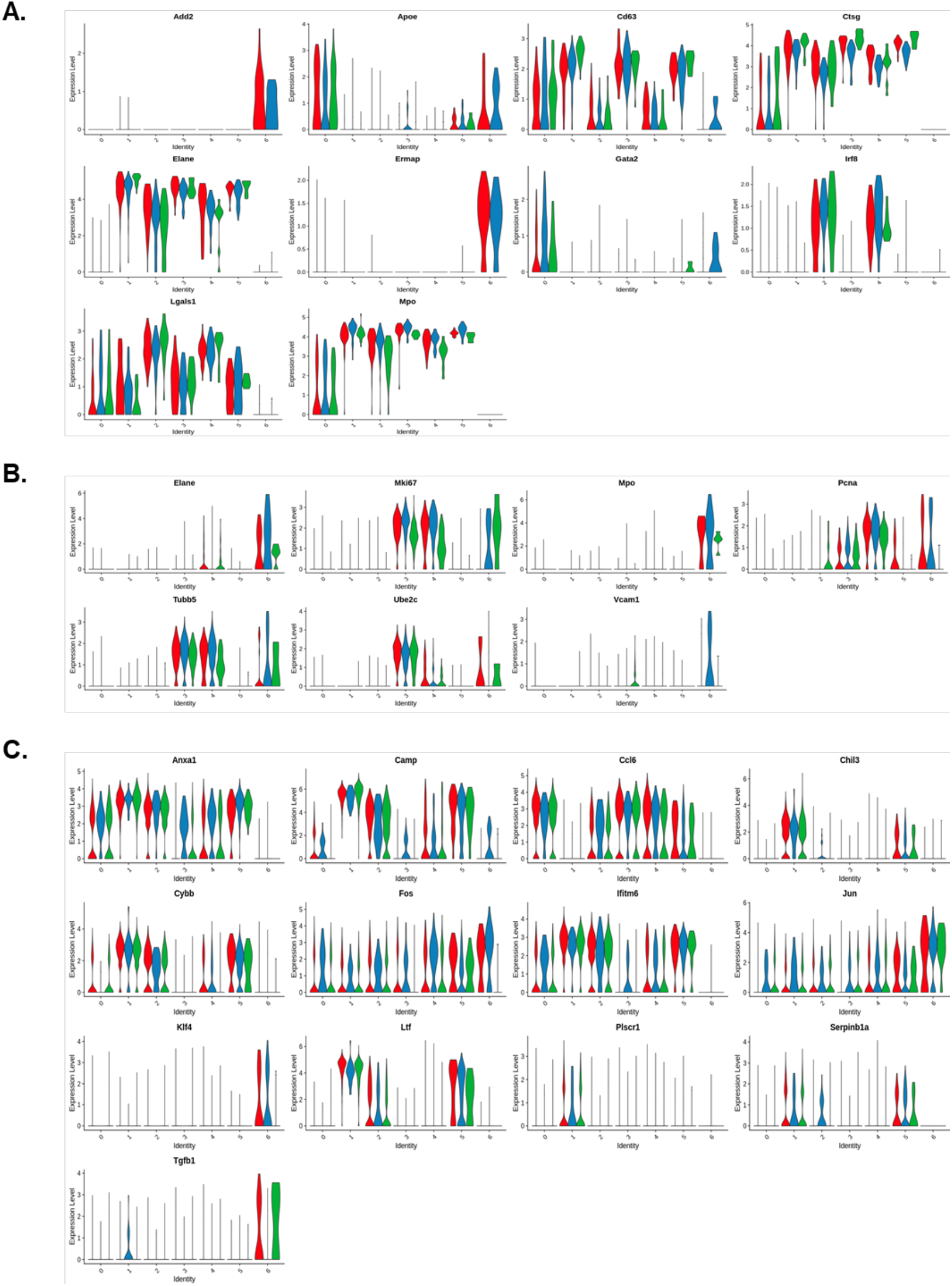
Violin plot of subcluster-specific gene expression. Red indicate young, blue indicate middle and green indicate old group.

**Supplementary Figure 7.**
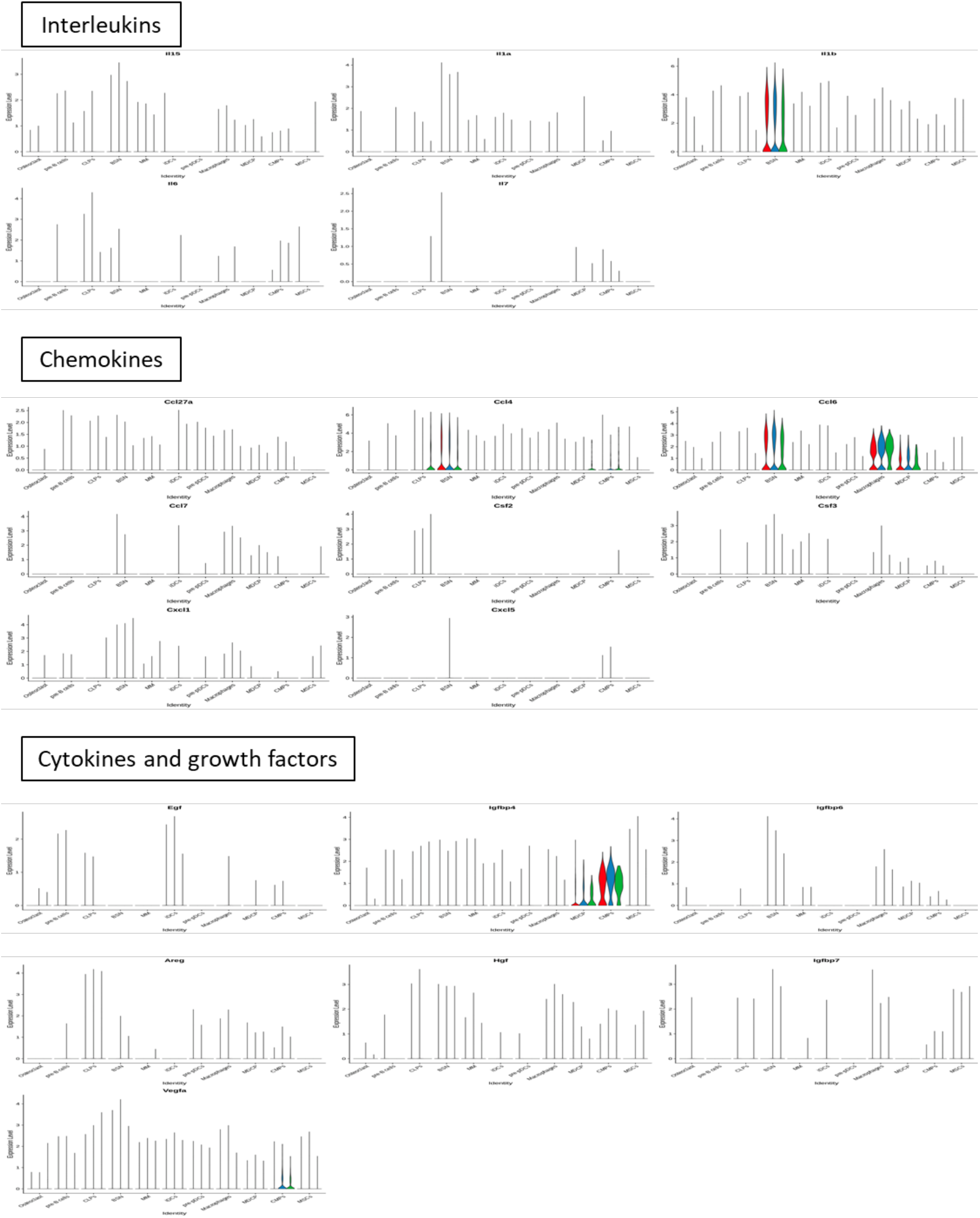
Cell type-specific secretory protein expression. Red indicate young, blue indicate middle and green indicate old group.

**Supplementary Figure 8.**
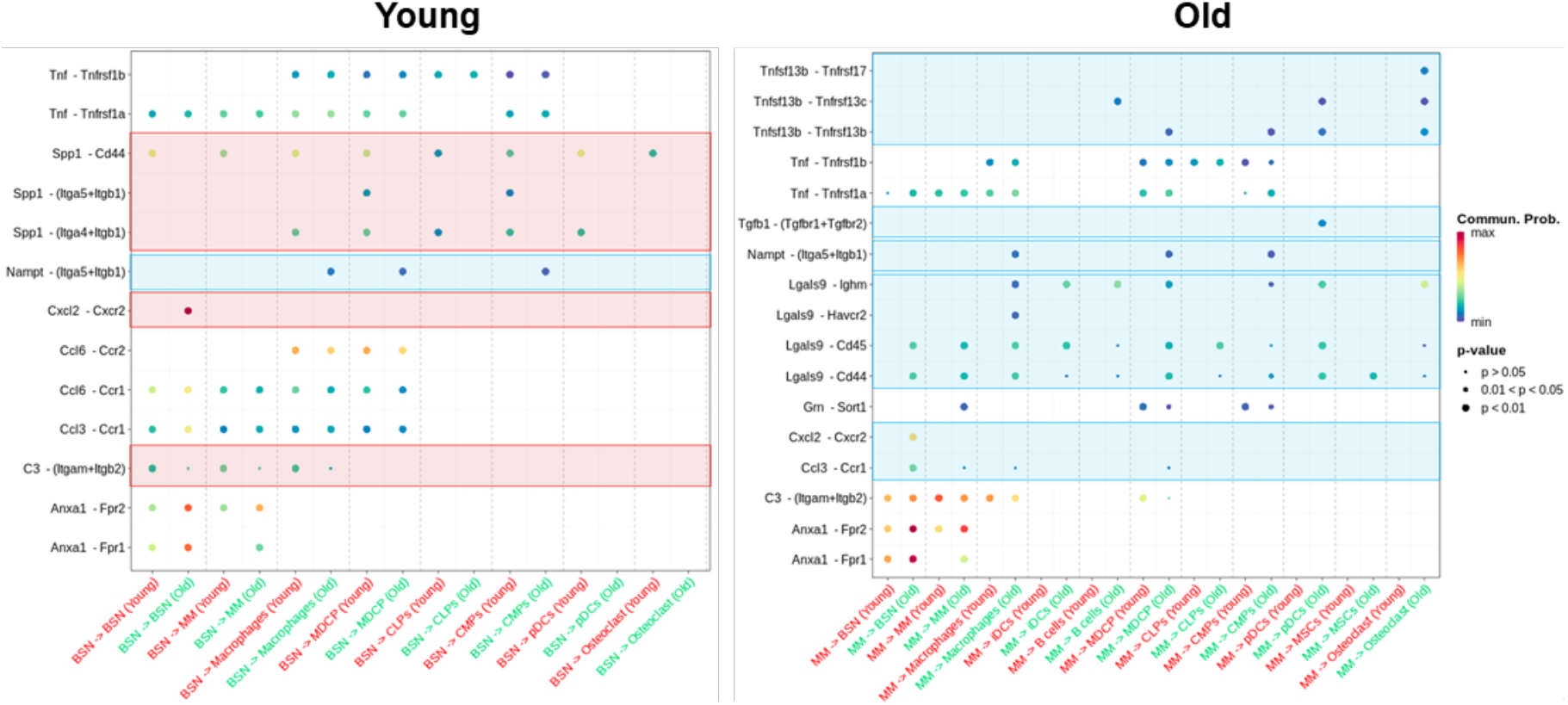
Age-associated ligand–receptor interactions in BSNs and MMs predicted by CellChat. The increased outgoing interactions from young BSNs to other cells marked red boxes and decreased interaction marked blue box (Young). The increased outgoing interaction from old MMs to other cells marked blue boxes (Old).

**Supplementary Figure 9.**
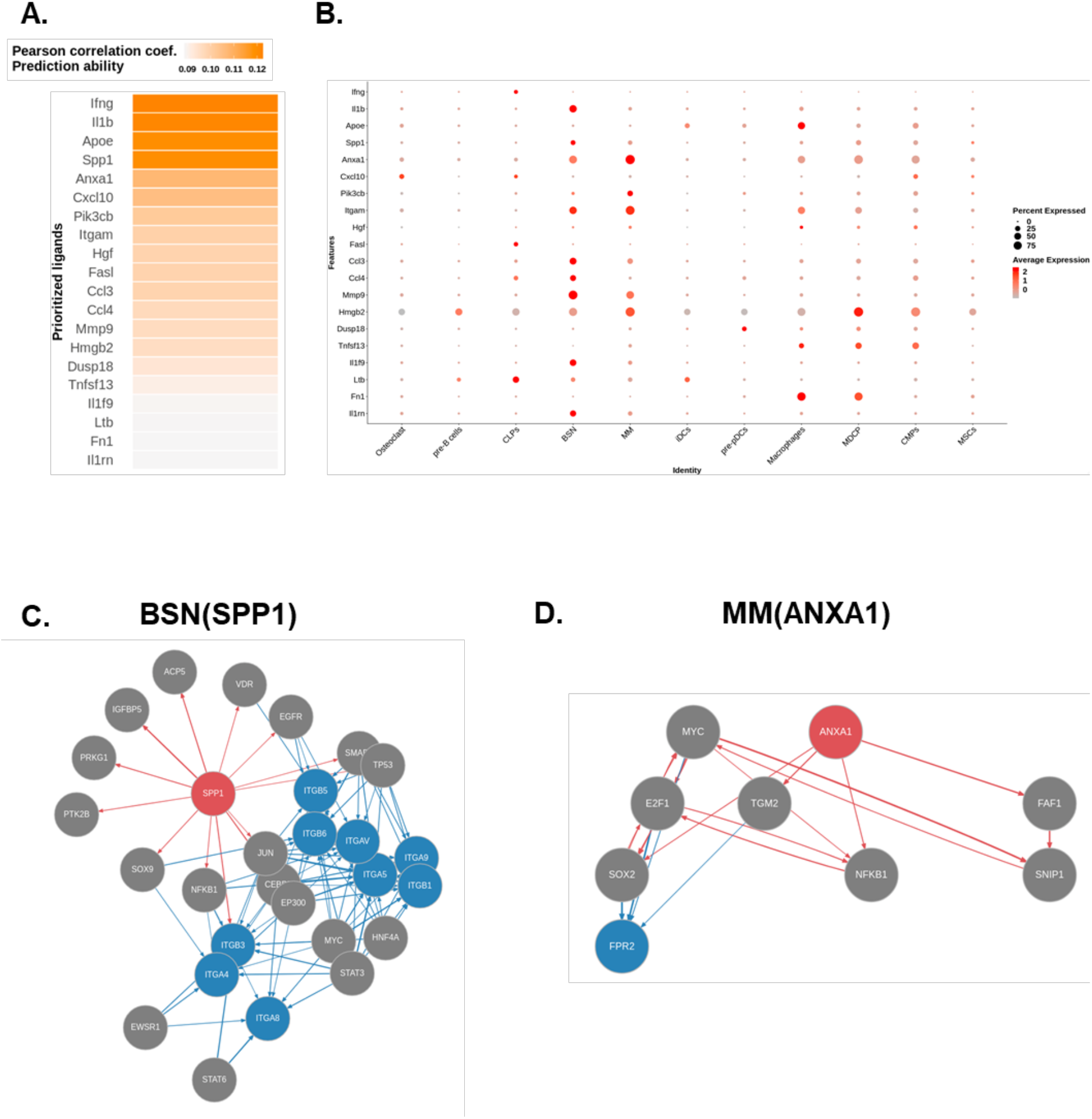
Significantly altered PPI in NicheNet analysis.

## Acknowledgements

This work was supported by Basic Science Research Program through the National Research Foundation of Korea (NRF) funded by the Ministry of Education (2017R1A4A1014584, 2020R1A4A1019423, 2017R1A2B3011778, and 2020R1A2B5B02002658 to H.M. R; 2021R1C1C2095130 to W.J.K; 2022R1I1A1A01062894 to K.T.K).

## Author contributions

Conceptualization was carried out by H.M.K., W.J.K. and K.T.K. K.T.K., J.I.M and S.G.P. were responsible for the methodology. Investigation was conducted by W.J.K. and K.T.K. Resources were the responsibility of W.J.K., K.T.K., J.I.M and S.G.P. Writing of the original draft was conducted by H.M.K., W.J.K. and K.T.K. Reviewing and editing was carried out by H.M.K., W.J.K. K.T.K., J.I.M., S.G.P., H.J.K., H.R.S. and H.I.Y. H.M.R. and Y.D.C. were responsible for supervision. H.M.R., W.J.K. and K.T.K. were responsible for funding acquisition.

